# A linguistic representation in the visual system underlies successful lipreading

**DOI:** 10.1101/2021.02.09.430299

**Authors:** Aaron R Nidiffer, Cody Zhewei Cao, Aisling O’Sullivan, Edmund C Lalor

## Abstract

There is considerable debate over how visual speech is processed in the absence of sound and whether neural activity supporting lipreading occurs in visual brain areas. Surprisingly, much of this ambiguity stems from a lack of behaviorally grounded neurophysiological findings. To address this, we conducted an experiment in which human observers rehearsed audiovisual speech for the purpose of lipreading silent versions during testing. Using a combination of computational modeling, electroencephalography, and simultaneously recorded behavior, we show that the visual system produces its own specialized representation of speech that is 1) well-described by categorical linguistic units (“visemes”) 2) dissociable from lip movements, and 3) predictive of lipreading ability. These findings contradict a long-held view that visual speech processing co-opts auditory cortex after early visual processing stages. Consistent with hierarchical accounts of visual and audiovisual speech perception, our findings show that visual cortex performs at least a basic level of linguistic processing.

## Introduction

The ability to exchange complex concepts and meaning through oral communication is a defining feature of human cognition. Although spoken language is primarily transmitted via acoustic signals, visible cues such as gestures and mouth movements are generated during speech production and play a prominent role in shaping our speech and language perception. In challenging listening environments when audible speech becomes contaminated by noise or other distractors, it is almost instinctual for listeners to look to the face for support. In doing so, both comprehension [1,2] and the ability to focus attention toward the speaker [3,4] are improved. These benefits are likely mediated in part by enhanced cortical tracking of speech acoustics [3,5] derived from correlated dynamics of mouth movements [6,7]. In contrast, visible speech cues can alter the phonemic perception of audible speech (i.e., the McGurk-McDonald Effect [8]) whereby incongruent audiovisual speech inputs (e.g., a visible /ga/ and an audible /ba/) combine to produce a percept that is absent in the speech signal (e.g., /da/), providing evidence for somewhat independent linguistic representations from acoustic and visible speech. Moreover, in other situations such as lip-reading, visible cues can form the entire basis of language comprehension [9].

Exactly how vision and the visual system contribute to the reception of spoken language, especially in very noisy conditions or in the absence of an acoustic cue, is a matter of debate. One view [10–14] which has garnered renewed attention holds that visual speech perception co-opts the specialized speech processing machinery of the auditory system and does so in early sensory processing stages (i.e., after extracting the temporal dynamics of lip movements). Recent accounts propose that auditory regions can “synthesize” a representation of the *unheard* acoustic speech envelope from the temporal dynamics of lip movements [15], which is possible due to the inherent correlation between acoustic and visible speech signals [6]. However, this view is complicated by two caveats. First, even when a speech signal can be heard, cortical envelope tracking does not guarantee comprehension. Although it does vary with intelligibility [16 18], envelope tracking has been observed during unintelligible speech [18] and non-speech signals [19]. Envelope tracking is also easily dissociable from comprehension when the envelope and temporal fine structure do not match (i.e., speech-speech chimeras [20]) and thus likely reflects general acoustic processing more so than speech processing. Second, the rise of this view, which was contemporaneous with our emerging understanding of broad and early multisensory interactions across neocortex [21,22], is consistent with the modulation of auditory cortical activity by generic (i.e., non-speech) signals from other modalities [23 25] and other non-specific multisensory enhancements bestowed by correlated temporal dynamics [26 31]. Therefore, it is unclear if visually-evoked envelope tracking in auditory cortex support language processing or if it simply reflects the dynamics of visual speech meant to enhance the processing of an acoustic signal that is typically present [7,32].

A separate view, supported by a trove of behavioral evidence, suggests that the visual system does its own heavy lifting during speech perception [33]. According to this perspective, each level of the speech hierarchy that is conveyed through acoustics are also available to the visual system and visual cortex contains the necessary machinery to extract and interpret the rich visible cues of speech. This view is further supported by two independent proposals. One [34] proposes that visual speech contains two distinct forms of information in relation to acoustic speech, viz. redundant and complementary information, and another [35] suggests that audiovisual speech processing occurs in a multistage process that includes early and late components. Redundant cues are those in which vision echoes the temporal dynamics of the articulatory patterns that form speech (i.e., correlated temporal information contained in lip movements, as discussed above) that are likely integrated in an early multisensory stage. In contrast, complementary cues supplement under-specified features of the acoustic speech and convey higher-level information to be integrated with acoustic cues in a late stage of integration. Phonemes that are easily confused acoustically can often be disambiguated by their visible articulations (e.g., /b/ and /d/, which begins to explain the McGurk-MacDonald effect). These complementary visual speech cues form - termed “visemes” - a compelling basis for lipreading [36,37]. It is plausible, if not likely, that visual cortex itself is performing some linguistic processing but there are few results in the literature that speak directly to how or even whether the different levels of speech that can be perceived visually [33], are represented in visual cortex. One study suggests that visual cortex does indeed track these so-called visemes in on-going speech [38] while another [39] provided evidence that visual cortex can easily discriminate perceptually-distant visemes, hinting at the idea that visual cortex does indeed perform some level of linguistic processing.

Recently, numerous studies have explored how the dynamics of silent visual speech are reflected in neural data [15,38 42]. In doing so, these studies have made great strides toward our understanding of the neural underpinnings of visual speech processing. However, their ability to speak to *linguistic* processing of visual speech comes with some important caveats. They have largely been limited by individuals’ naturally poor lipreading skill, which limits the extent of measurable linguistic activity in the brain; they have often lacked critical behavioral measures, limiting the ability to relate neural activity to language reception, which is highly variable across individuals [33]; and perhaps most importantly they often utilize an impoverished proxy measure of visual speech (e.g., the acoustic speech envelope), which limits the potential to disentangle contributions of different hierarchical processing stages during the processing of visual speech. In the current experiment, we address these caveats to probe whether linguistic processing of visual speech occurs in visual cortex. To this end, we asked human observers to rehearse a set of audiovisual videos of a speaker with the intent of lipreading them later. We then recorded EEG from the participants as we replayed silent versions of the rehearsed videos and new set of videos without sound. Finally, we asked participants to detect target words during the silent videos and asked them to subjectively rate their intelligibility. In our paradigm, lipreading ability was improved by the rehearsal. We find neurophysiological evidence that speech-specific (lip movements) and language-specific (visemes) visual features were represented in separable components of ongoing brain activity. Of all the features we tested, only the linguistic feature was enhanced by the rehearsal and that enhancement was reflected across participants lipreading scores.

## Results

### Occipital electrode recordings reflect the encoding of linguistic features during silent speech

We recorded 128-channel EEG from sixteen participants as they watched new and previously rehearsed silent videos of continuous natural speech of a male speaker discussing political and economic topics (Figure 1a). To relate their EEG responses to the speech, we first extracted several features of the visual stimulus (Figure 1b) including motion flow (M), vertical and horizontal lip movements (L), and a set of 12 visemes (V). Using linear modeling, we then derived a temporal filter relating each stimulus feature to the low-frequency EEG (Figure 1c). This filter the temporal response function (TRF) describes how fluctuations of a stimulus feature impacts neural activity across a set of time lags and can be envisioned as analogous to the more conventional event related potential (ERP). Using a cross-validated approach, we quantified how well each feature was represented in the neural data, aiming to directly compare the quality of each representation. We fit a TRF using a subset of trials and then predicted EEG data from left out trials by passing the stimulus features of that trial through the TRF. We measured the accuracy of predictions, channel-by-channel, by calculating Pearson’ correlation between the real EEG and the predicted EEG signals.

**Figure 1.**
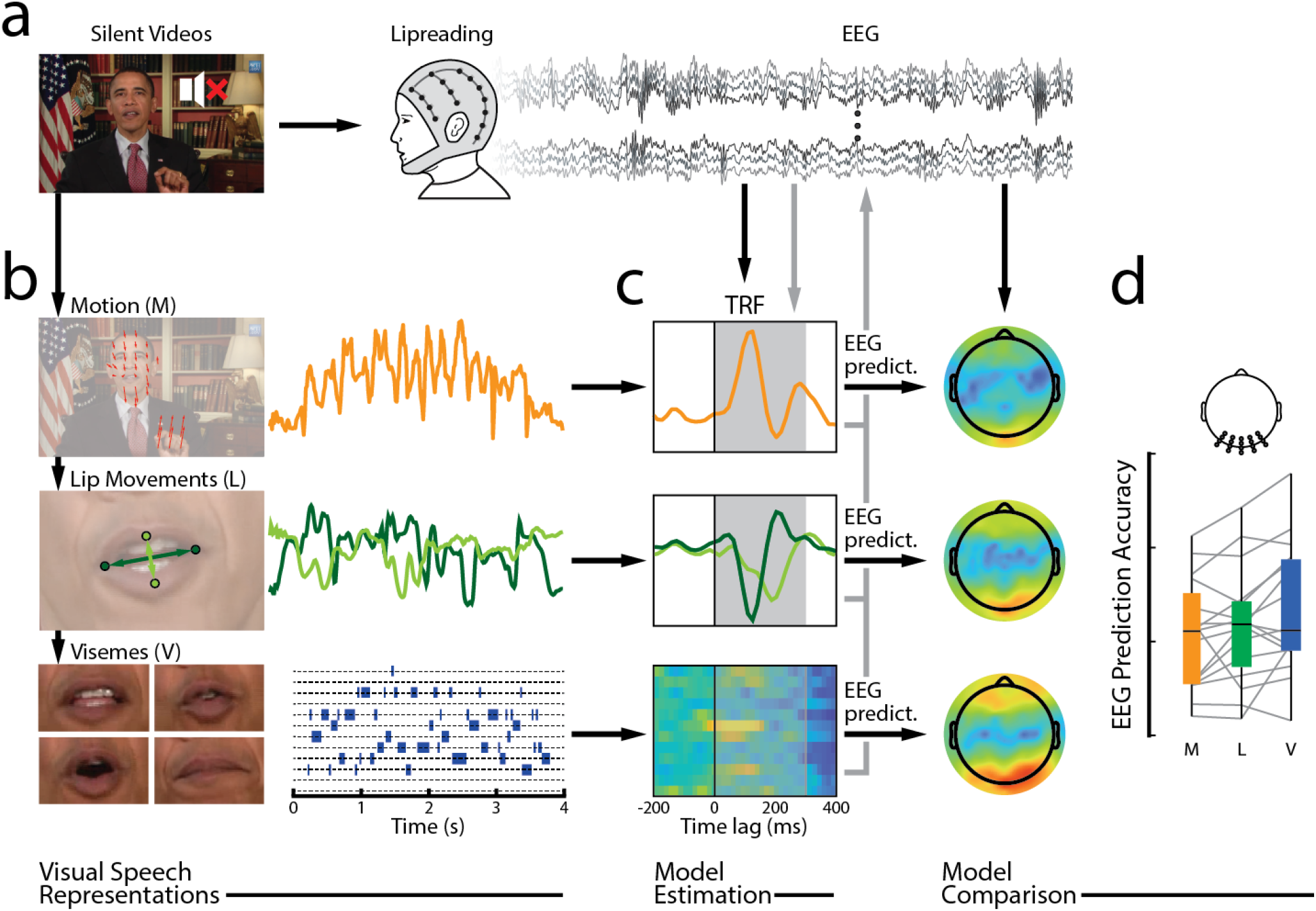
| Modeling different features of visual speech. **a**. We recorded EEG signals while participants watched minute-long videos of silent speech and attempted to lipread. **b**. From the videos we extracted several features to serve as regressors in our TRF analysis. **c**. An extracted feature, or a combination, was regressed against EEG activity across a series of time lags on a subset of data. The resulting TRFs (average across the twenty occipital scalp electrodes shown here) could then be used to predict EEG on left-out data. These predictions could be used to evaluate the models (black arrows) or could be subtracted from the original EEG to remove contributions of that feature (grey arrows) before predicting other features. To standardize the predictions across features, we used the same time lags (0 300 ms, grey boxes) for each model prediction. **d**. To compare across models, we calculated the average prediction accuracy across a set of 20 occipital electrodes.

Our first analysis sought to replicate and extend a previous finding [38] demonstrating that visemes are represented in neural activity. Focusing our analysis on 20 electrodes over occipital scalp (Figure 1d), we quantified the ability of each visual feature to predict EEG activity recorded while participants viewed five *previously unseen* silent videos. By comparing these predictions to a permuted null distribution, we found that neural activity in occipital electrodes could be reliably predicted by models derived visemes (V; T(15) = 2.94, p = 0.010), lip movements (L; T(15) = 4.78, p = 0.00024, and visual motion flow (M; T(15) = 4.37, p = 0.00054).

An important consideration in this analysis is that these features share some temporal structure (i.e., they are correlated). So, to better disentangle the contribution of visemic encoding in neural activity, we took a bi-faceted approach. First, in line with similar work in the acoustic domain [43] we constructed a family of models consisting of combinations of each feature. The logic behind this approach suggests that if a joint-feature model (e.g., ML) out-performs its constituent single-feature model (M or L), then EEG indexes the processing of both features. Indeed, we found that when adding V to either low-level feature (M or L) or their combination (ML), model performance improved (Figure 2a; MV vs. M: T(15) = 4.26, p = 0.00068; LV vs. L: T(15) = 4.55, p = 0.00038; MLV vs. ML: T(15) = 4.84, p = 0.00022). Second, we utilized the predictive power of our TRF analysis to construct and remove predictions of EEG activity related to lip movements and general motion flow from the EEG signal [after 44]. This approach effectively eliminates the EEG signature of the partialed-out feature (see Methods and Table 2). We then predicted the residual EEG using visemes (Figure 2c). The performance was unsurprisingly lower in these reduced models due to the removal of the covariant activity yet remained reliably better than shuffled permutations when removing motion flow (T(15) = 2.65, p = 0.018), lip movements (T(15) = 2.44, p = 0.028), or both (T(15) = 2.15, p = 0.048). Taken together, these results give support to the idea that visual cortex is capable of modest linguistic processing even with minimal language comprehension.

**Figure 2.**
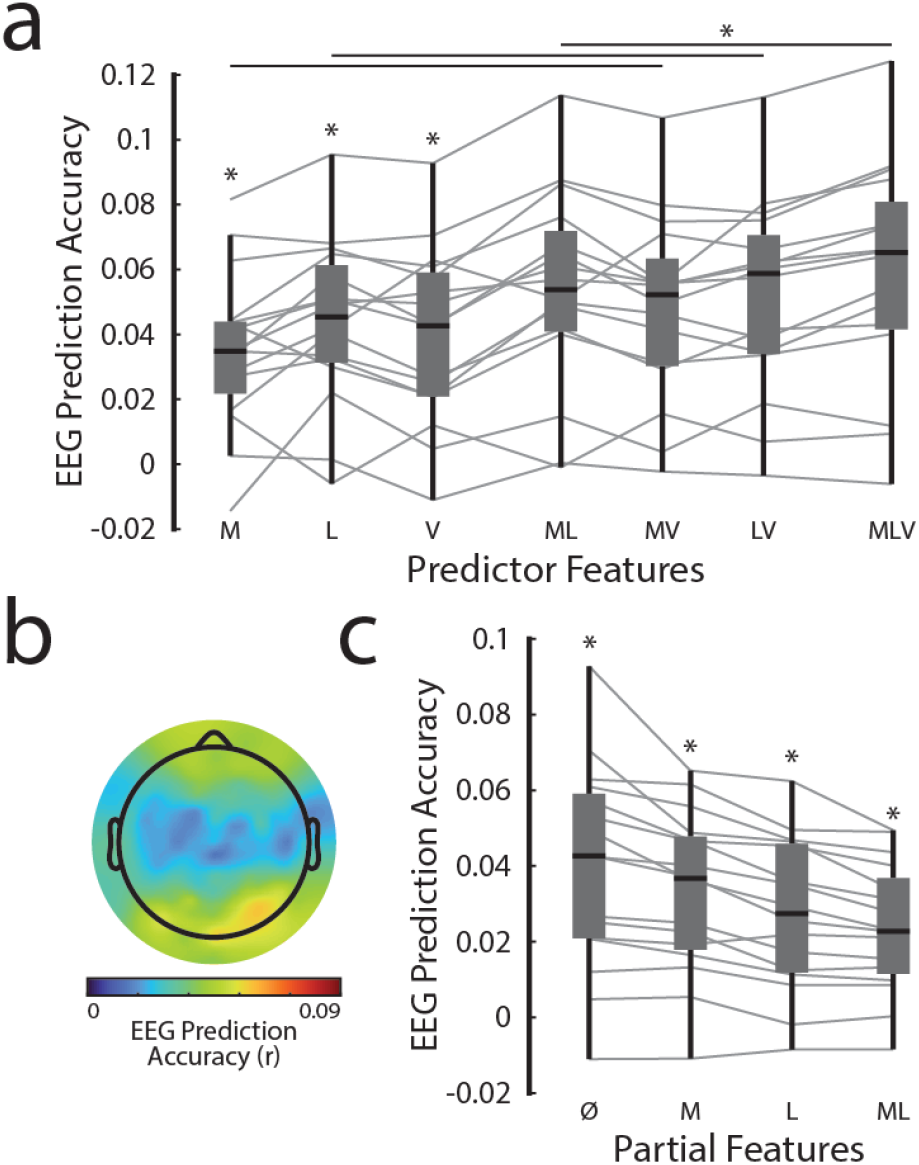
| Visual responses recorded while participants viewed novel videos reveal the brain’s representation of visemes. **a**. Comparison of prediction accuracy from the full family of models during novel video presentations. Notably, adding visemes (V) to any model (M, L, or ML) improved EEG predictions, suggesting separable contributions of visemes and the low-level features in brain activity. **b**. Topographic representation reveal a robust representation of visemes in EEG activity recorded over occipital scalp. The color bar shows the scale of all topographic plots in this article **c**. Partialing motion (M) and/or lip movements (L) reduced the strength of viseme-based predictions, yet those predictions remained significantly above chance indicating robust viseme tracking.

### Successful lipreading reflects improved visual linguistic processing

Prior to the EEG session, we asked participants to mentally rehearse five videos with sound with the goal of being able to later identify target words based on tracking the speaker’s visible speech articulations. During the EEG session, participants viewed silent versions of these rehearsed videos along with the previously described unseen videos. During each trial, participants detected target words given at the start of the trial and gave a subjective intelligibility rating of that trial at its conclusion. Both the objective performance (d’; T(15) = 3.45, p = 0.0036) and the subjective intelligibility rating (T(15) = 3.27, p = 0.005) were higher for rehearsed videos suggesting that participants were able to better lip-read those videos (Figure 3).

**Figure 3.**
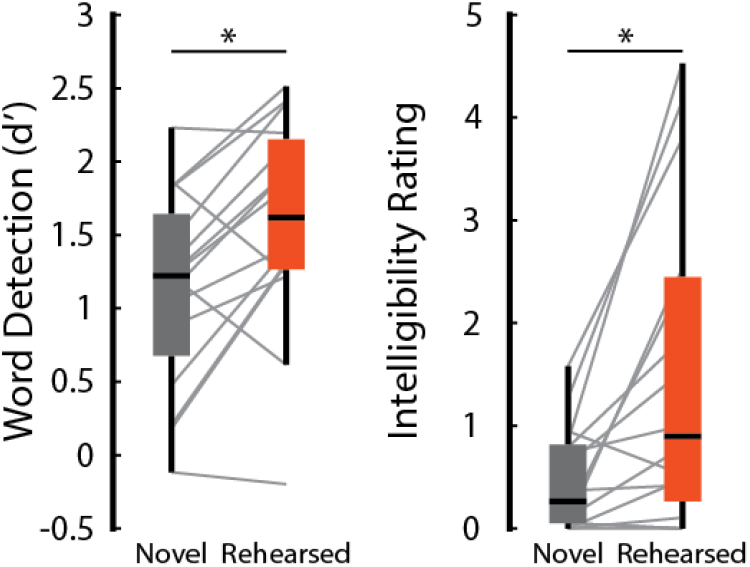
| Speech rehearsal improved lipreading and subjective intelligibility during silent speech. During the silent videos, participants were asked to report their detection of a target word with a button press. After each video, they also rated the subjective intelligibility of the speech. Rehearsal improved both measures across participants.

With this evidence that our rehearsal paradigm was behaviorally successful, we next turned our attention toward the neural underpinnings of participants’ improved ability to read lips. First, we compared the prediction accuracy of a family of models across features and rehearsal conditions. We performed a two-way, repeated measures analysis of variance (ANOVA) which found significant main effects for both feature (F(2,30) = 8.98, p = 0.00088) and rehearsal condition (F(1,15) = 4.80, p = 0.045). Importantly, this analysis also found a significant interaction (F(2,30) = 4.82, p = 0.015), suggesting that rehearsal might be preferentially affecting the encoding of certain features. Follow-up pairwise comparisons showed that this was indeed the case (Figure 4a): viseme encoding improved with rehearsal (T(15) = 2.96, p = 0.0097) as did motion flow (T(15) = 2.17, p = 0.046), but lip movement encoding did not(T(15) = 1.08, p = 0.29). Viseme encoding was stronger than motion flow (T(15) = 3.75, p = 0.0020) and lip movements (T(15) = 4.93, p = 0.00033) during the rehearsed videos while visemes (T(15) = 2.18, p = 0.046) and lip movements (T(15) = 2.17, p = 0.047) were better predictors of EEG than motion flow during novel videos. These results suggest that rehearsal confers an advantage to the encoding of linguistic features (visemes) but not speech features (lip movements).

**Figure 4.**
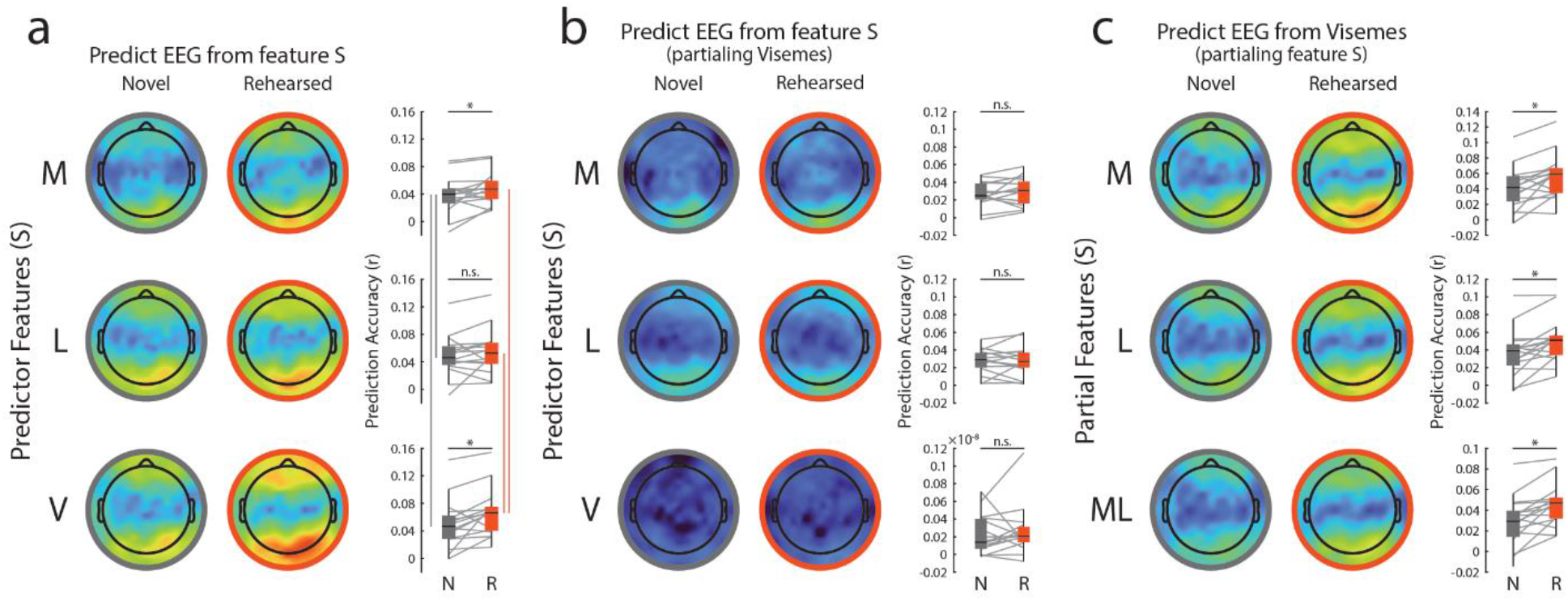
| Visemes are preferentially improved after lip-reading rehearsal. **a**. Topographic representation of the scalp show EEG prediction accuracy (left) when predicting from each stimulus feature (S = motion [M], lip movements [L], or visemes [V]) during novel (grey) and rehearsed (red) conditions. Box and whisker plots (right) of the average prediction accuracy over a set of 20 occipital electrodes show that rehearsal preferentially improved the representation of visemes. **b**. Topographic representation of each stimulus features (S = M, L, or V) after partialing visemes from EEG show the contributions of motion flow and lip movements separate from linguistic units. Importantly, we were able to observe residual responses to both low-level features, and neither showed any rehearsal-based enhancement over occipital scalp. Note the drastic reduction of model performance (∼8 orders of magnitude) when predicting visemes from “viseme-free” EEG (bottom; see also Table 2). **c**. Topographic representation (left) of the viseme predictions after partialing S (S = M, L, or ML). Box and whisker plots of the average prediction over occipital scalp show that viseme predictions are enhanced by rehearsal even after partialing contributions of other visual features.

As before, we took a complementary and more direct approach to unravel the neural consequence of rehearsal involving isolating the contribution of a feature of interest by regressing out the contributions of other features. These analyses were necessary to account for correlated structure and shared predictive power between the features and to clarify ambiguous findings related to how motion flow encoding varies with rehearsal. As expected, there were no rehearsal-based improvements to neural tracking of motion flow (T(15) = 1.14, p = 0.27) or lip movements (T(15) = 0.15, p = 0.90) after regressing out the contributions of visemes (Figure 4b). But importantly both features were still robustly encoded above chance during both rehearsed (M vs shuffled M: T(15) = 5.30, p = 0.000089; L vs shuffled L: T(15) = 4.80, p = 0.00024) and previously unseen videos (M vs shuffled M: T(15) = 4.76, p = 0.00026; L vs shuffled L: T(15) = 5.31, p = 0.000088). Conversely, the benefit of rehearsal on viseme tracking remained intact (Figure 4c) after partialing out motion flow (T(15) = 3.17, p = 0.0064) and lip movements (T(15) = 3.24, p = 0.0055) separately and in combination (T(15) = 3.47, p = 0.0034). These analyses suggest that although there is likely some neural overlap of the processing of visual speech features and visual language, we are able to index them separately and observe that linguistic processing is preferentially improved when participants can lipread silent speech.

The previous analyses identify viseme tracking over occipital scalp as the likely driver of lipreading ability. To explore this possibility more directly, we examined behavioral scores across participants in the context of the neural signature of viseme processing. Specifically, we quantified improvement in the word detection task and improvements in viseme tracking and tested their relationship. This comparison revealed a robust positive correlation (Figure 5a; r = 0.60, p = 0.015). This relationship was not abolished after partialing motion flow (r = 0.57, p = 0.021), lip movements (r = 0.55, p = 0.028), or both (r = 0.52, p = 0.040) from the EEG. Thus, we take this as evidence that neural activity specific to the processing of visual linguistic information is crucial to supporting lipreading.

**Figure 5.**
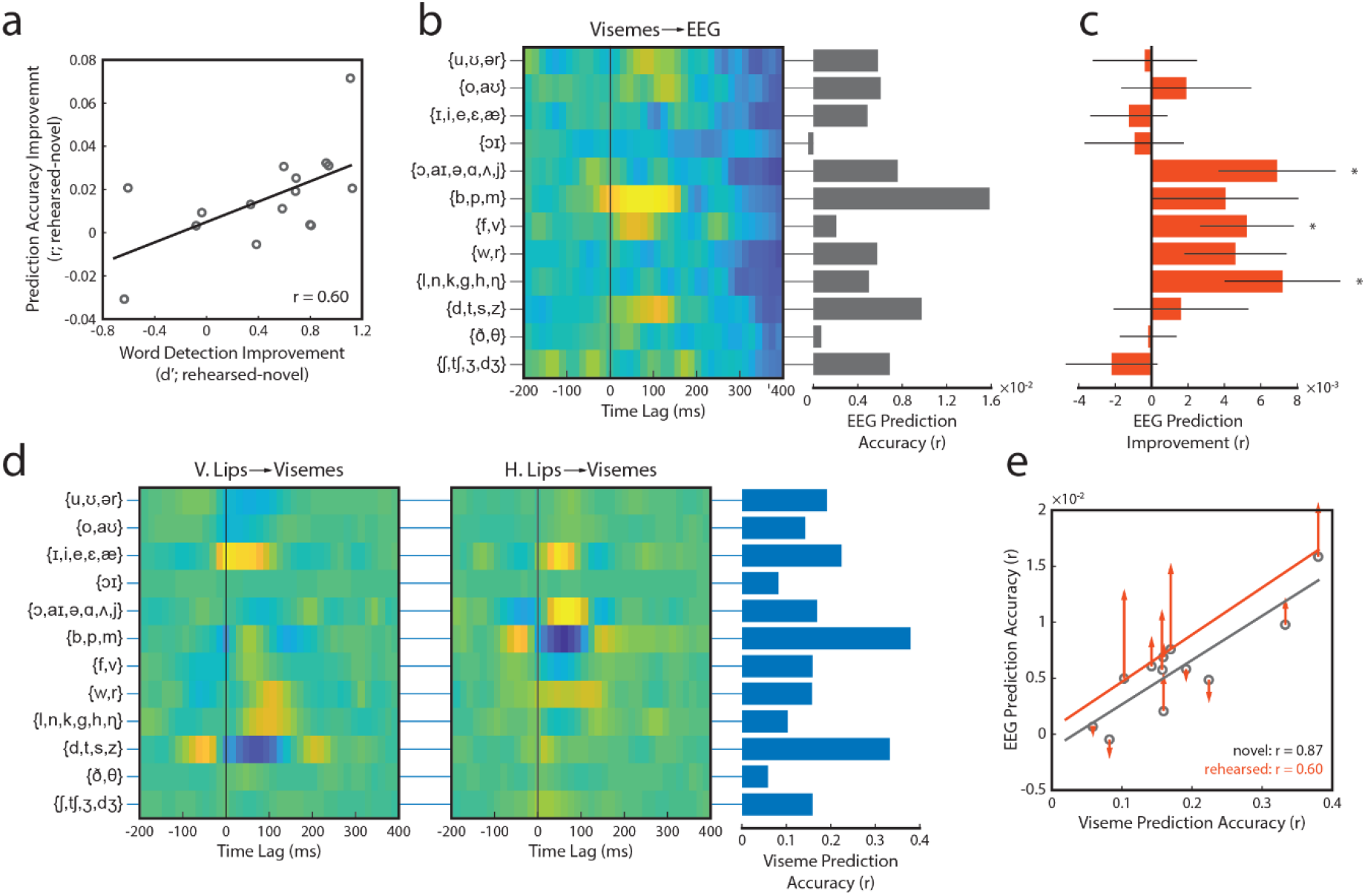
| Individual viseme analysis. **a**. Across participants, the viseme-based EEG prediction and lipreading improvements related to rehearsal were correlated. **b**. We fit a multivariate TRF that predicts EEG activity from visemes (left) before separating into individual viseme TRFs and predicting EEG with each. With those univariate TRFs, we predicted left-out EEG data and measured their individual prediction accuracies (right). Shown is prediction accuracy during novel videos. Visemes vary in their individual contribution to the EEG signal. **c**. Individual viseme encoding improvements conferred by rehearsal. Darker colored bars represent the average improvement across participants and conditions. Three visemes were significantly better EEG predictors after rehearsal. Black line and light shaded region represent the expected improvements based on shuffled permutations. **d**. We fit TRFs to predict visemes from vertical (left) and horizontal (middle) lip movements. Here, the TRF represents the average lip movements for each viseme. The ability of lip movements to predict visemes (right) were variable across visemes in a manner similar to (**b**). Dark colored bars represent the average across stimuli while black lines and light shaded regions represent the mean and standard error of shuffled permutations of these predictions. **e**. The extent that each viseme is encoded in brain activity is largely predicted by how well it is specified by lip movements, especially during novel videos (black line). This relationship is disrupted somewhat by rehearsal (red line), owing to changes (red arrows) that were unrelated to the lip-viseme relationship described in (d).

### Specific individual visemes contribute to improved linguistic processing in visual responses

Having established a connection between lipreading and the neural activity that tracks visemes in ongoing speech, we undertook an exploratory analysis to unravel contributions of individual viseme categories. We predicted EEG activity from individual (i.e., univariate) viseme models and evaluated the strength of those predictions. To account for dependencies between visemes, we first fit a multivariate viseme TRF (as we’ve done for all viseme analyses up to this point), then separated that model into individual univariate TRFs [after 45], and finally predicted left-out EEG data with each univariate model (Figure 5b, novel shown). We submitted the resultant prediction accuracies to a repeated-measures ANOVA (with factors viseme category and rehearsal condition). A significant main effect of viseme category indicated that individual visemes do not contribute equally to the EEG signal (F(11,165) = 11.00, p = 4.1×10^−15^) with some being more strongly encoded than others. We wondered if this differential encoding of visemes may be indicative of a mapping between lip movements and visual linguistic units that participants could be reinforcing during the rehearsal. That is, we sought to reveal any potential relationship between lip movements and visemes and test whether rehearsal reinforced that representation in brain activity. We fit TRFs that predict visemes from horizontal and vertical lip movements and found that like EEG predictions from visemes, visemes are differentially predicted on the basis of lip movements (Figure 5d; F(10,140) = 90.01, p = 4.69×10^−56^). Note that we removed the fourth viseme ({ɔɪ}) in this analysis due to its under-representation in four of the fifteen video clips. In a complementary analysis, we removed those four video clips from the analysis (keeping all visemes), which yielded the same result (F(11,110) = 68.52, p = 3.66×10^−44^). In fact, the degree to which EEG encodes each viseme individually is highly correlated to how well those visemes are described by lip movements (Figure 5e, grey line; r = 0.87, p = 2.04×10^−4^) for the unrehearsed videos, suggesting that linguistic processing in the absence of successful lip reading may be largely attributable to how well lip movements are encoded in brain activity. However, when observers can successfully lipread, this relationship is perturbed slightly (Figure 5e, red line; r = 0.60, p = 0.019). Importantly, this interruption is driven by the enhancement of individual visemes in a manner that is unrelated to the lip-viseme mapping (Figure 5c and Figure 5e, red arrows; r = 0.07, p = 0.82), suggesting rehearsal was not reinforcing this mapping. Specifically, three viseme categories ({ɔ,aɪ,ə,ɑ,ʌ,j}, {f,v}, and {l,n,k,g,h,η}) drive this enhancement derived from rehearsal (T(15) = 2.50, 2.38, and 2.64; p = 0.025, 0.031, and 0.019, respectively). Taken together, these exploratory analyses suggest that participants do not simply learn to categorize lip *movements* when they learn to read lips. Instead, it seems more likely that rehearsal-based improvements rely more on lip *shape*, as suggested by Campbell [34]. Future studies with better representation of individual visemes and more powerful analyses will be needed to confirm these findings.

## Discussion

In the current study, we recorded brain activity over occipital scalp during silent speech and related it to a set of visual features derived from the speech signal. We were able to isolate the activity related to linguistic features from activity related to lip movements and other motion present in the videos. In doing so, we show that this linguistic representation is robustly tracked in ongoing brain activity, replicating and extending previous work from our group [38]. We found that brief, directed rehearsal of speech promotes an improvement in lipreading and preferentially improves the representation of visual linguistic features of that speech. Finally, we linked behavioral improvements across participants to stronger tracking of linguistic content in visual brain regions, contrary to the suggestion that lipreading is supported by activity in auditory cortex [10 15].

Recently, a flurry of studies have made substantial contributions to our understanding of the neural processing of visual speech and its contributions to audiovisual speech processing [15,38 42]. Although some provide promising evidence of specialized speech processing in visual cortex [38,40,41], none have yet to provide a link between *meaningful* reception (i.e., lipreading) and visual brain activity. For example, several studies have related the tracking of an unheard acoustic envelope to phonological processing [15,40], though even during acoustic speech recpetion the specificity of envelope tracking to linguistic processing is questionable. Neural signals track the envelope of both non-speech signals [19] and unintelligible speech [18,46]. Envelope tracking does not necessarily index anything beyond energetic fluctuations [20] since the broadband amplitude envelope derived from speech signals does not provide enough information to support phonological processing or comprehension [18,47]. Due to shared dynamics between acoustic envelope and lip movements [6], these visual speech studies are potentially indexing activity related to lip movements, which is not affected by lipreading according to our findings. One study contrasted forward and reversed visual speech as a proxy for comprehension [40] although it is unclear if there is any meaningful intelligibility to be lost by reversing *visual* speech or if neural differences could be explained by the unnatural temporal dynamics of reversed movement. Another study found that responses to visual speech were enhanced in mouth-responsive regions in visual cortex [41], yet these enhancements likely represent attentional preference to task- and context-relevant signals rather than linguistic processing. Finally, one study indexed linguistic processing directly [38], but provided no measurement of speech reception. Here we replicate and further validate this previous work from our lab by introducing a necessary control variable related to lip movements and extra analytical controls.

Across each of these studies, the meaningfulness of the reception is uncertain given the lack of behavioral measurement and when considering how poor and variable human lipreading is [48,49]. Our approach brings a crucial behavioral measure that was lacking in this literature. By relating our neural findings to simultaneously acquired behavior, we find strong evidence to support a basic level of linguistic processing in visual cortex. Previous investigators have suggested that a novel area in visual cortex, which was given the moniker temporal visual speech area (TVSA), exhibits activity patterns suggestive of specialized speech processing [50]. Using contrasts across several stimulus conditions, they found no support for specialized speech processing in other brain areas engaged in processing visual motion (e.g., middle temporal visual area) or face (e.g., fusiform face area) that have been reported to be active during visible speech [33,51]. We are limited by the (lack of) spatial resolution of our approach and cannot precisely localize visemic processing to specific visual regions. However, we were able to dissociate the neural signature of visual linguistic processing from activity related to lip movements themselves (Figure 4b,c), consistent with findings from Bernstein et al [50], although their claim stops short of crediting TVSA with visual speech categorization duties. It seems possible that TVSA is indeed involved in this process. It is also very probable that this process requires coordination from higher-order brain regions [40]. Regardless, some form of visual linguistic categorization is evident within visual brain areas.

Our findings also corroborate several findings and propositions related to audiovisual speech. Visible speech cues are important in the context of acoustic speech processing and have been described as providing two types of information redundant and complementary during audiovisual speech integration [34]. Redundant information refers to the correlated dynamics of visual and acoustic speech [6]. Visual timing information can “restore” their degraded acoustic speech correlate [7,27]. However, the benefits of correlated visual cues aren’t restricted to speech processing [52,53], they also encourage perceptual object formation [54 57], and promote attentional selection [3,4,28,58]. These effects have been demonstrated in low-level sensory cortex [59]. Therefore, it is likely that visual dynamics do not provide any linguistic information on their own but need to be transformed into such.It’s becoming clear from the current findings and others [38,39] that visual cortex performs that transformation, encoding a representation of visual speech that fulfills Campbell’s *complementary mode*. Complementary visual information refers to cues that are underspecified in the acoustic stream but conveyed more effectively in visible signals. For example, some speech units (e.g., /b/ and /d/ or /m/ and /n/) can be acoustically ambiguous, especially in noisy conditions, but their visible articulatory patterns are distinct and robust to noise. Although there is yet to be direct evidence for the integration of viseme and phoneme cues, indirectly the enhancement of acoustic representations (i.e., spectrogram) is similar in noisy and noise-free conditions while the enhancement of phonetic information appears to be more pronounced in noisy conditions [60], consistent with the role of complementary visual cues.

The processing of acoustic speech has been shown to be hierarchically organized, with distinct neural signatures of different levels of the linguistic hierarchy [43,61 63]. Our findings of separable visual speech and linguistic components suggest that visual speech perception is similarly hierarchical, at least up to the basic linguistic units. Together, these findings support the concept of a hierarchical structure of audiovisual speech integration [34,35]. Early integration and binding [56] may take the form of redundant visual cues modulating acoustic processing through direct connections between early visual and auditory cortices [64 66] while late integration may be related to the perceptual system deriving a language representation by drawing from independent and complementary acoustic- and visual-based linguistic signals. Further, these dual roles of visual speech may explain the concept of the “sweet spot” for audiovisual speech integration [48]. As speech becomes contaminated by low to moderate levels of noise, redundant visual timing cues are able to “restore” features in the acoustic stream [7] in tandem with an increase weighting of the visual speech representation [67], leading to a peak in visual enhancement of acoustic speech at a signal-to-noise ratio of about −12 dB. But as noise levels increase further, the acoustic stream may become too corrupted for visual timing cues to repair them and observers are forced to rely exclusively on visual speech cues.

There are a couple of caveats to our findings that are worth noting. First, it is possible that attention could explain the improvement in our neural measures [68]. According to this explanation, participants would be more engaged with a speech video that they were familiar with. Because of the nature of our rehearsal paradigm, it is impossible to monitor for this effect in behavior. We therefore performed control analyses to exclude the possibility of an attentional effect. If participants were attending more to the rehearsed videos, we would expect improved tracking of each of our visual features in the rehearsed condition [56,69,70]. Yet, even after we leveraged the predictive ability of our modeling framework to isolate activity specific to each feature, we found no evidence of improved representation of a general stimulus feature (motion flow) or a more specified task-relevant feature (lip movements). Any rehearsal-related improvements were consistently restricted to linguistic representations. We therefore think attentional effects are unlikely to be driving our findings. Second, it is possible that we are under-characterizing the non-linguistic speech features in our study. We made a decision to capture the dynamics information related to lip movements than some previous endeavors (i.e., by measuring horizontal and vertical dimensions [e.g., 15] rather than a one-dimensional measure of mouth area [e.g., 6,42,58]) due to the correlation between certain dimensions to distinct acoustic features (e.g., horizontal expansion and contraction co-varying with phonetically-relevant spectral cues [7]). It is still possible that a more complete characterization of the articulators (e.g., tongue movements) may account for some of the performance benefit of our viseme model. Since tongue movements are manifested in speech acoustics, characterizing their dynamics is surely to be informative on the nature of redundant multisensory interactions. But because the identity of any particular viseme does not depend on the visibility of the tongue, we think characterizing its movements is unlikely to account for the activity related to our viseme model or its improvement derived from lipreading. By their very nature, complementary visual cues such as visemes describe features orthogonal to speech dynamics, so features that de of speech dynamics or measure that is a derivative thereof [after 71] is unlikely to fully account for effects ascribed to complementary cues. Instead, a principled and stimulus-based description of the shape of the articulators is more likely to be fruitful in this endeavor.

## Conclusions

Here, we have provided evidence for specialized categorical linguistic processing in the visual system. This idea is strongly supported by multitude of behavioral observations suggesting a visual linguistic structure that reflects its acoustic counterpart [33] but with its own idiosyncratic structure that is a consequence of phoneme confusion [72,73]. Such a correspondence between auditory and visual speech processing hierarchies, which itself is a reflection of the common source of acoustic and visual speech cues, is essential to multistage integration of audiovisual speech [35]. In this multistage integration, low-level visual cues (such as lip movement dynamics) do have an entry into early auditory cortex [e.g., 15], but this convergence of *redundant* visual cues likely serves more general, non-linguistic processes that benefit speech perception: to increase auditory cortical sensitivity to upcoming events [74], to perceptually restore degraded acoustics [7,27], and to facilitate attentional selection of auditory signals [3,4,28]. The higher-order complementary linguistic representation demonstrated here is surely a major contributor to the late integration stage of Peelle’s model where it would be aptly suited to constrain the inference of the identity of a spoken utterance. Finally, we find evidence that a rudimentary lipreading ability is supported by the neural tracking of visemes. Thus, this neural activity may be the brick and mortar of visual language. It will be interesting for future work to shed light on how the visual system builds upon these basic linguistic representations and how far the sophistication of spoken language processing in the visual system extends. We also look forward to understanding the similarities and differences of visual linguistic processing in lipreaders who are deaf or hard of hearing.

## Methods

### Participants

Sixteen individuals (age = 25 ± 6.3, 9 females) participated in the current study. All participants reported normal or corrected-to-normal vision and normal hearing and were right-handed. The study was conducted in accordance with procedures approved by the University of Rochester Human Subjects Review Board. Informed written consent was obtained from all participants prior to any procedures. When applicable, participants were given monetary compensation for participation.

### Apparatus and Stimuli

The stimuli used in the current experiment consisted of videos of a well-known male speaker. The videos contained the speaker’s head, shoulders, and chest centered in the frame.The speech was conversational and directed at the camera, and the linguistic content focused on political policy. Fifteen 60-s videos were rendered into 1280 x 720-pixel movies in VideoPad Video Editor (NCH Software) at 30 frames-per-second. Audio was digitized at 48000 kHz with 16-bit resolution. A very brief (10 ms) audible click was inserted at the beginning of the audio track for EEG synchronization (see below).

Videos were presented via Presentation Software (Neurobehavioral Systems) on a desktop computer running Windows 10. Visual stimuli were presented on a 24” widescreen (16:9) LCD monitor (BenQ ZOWIE XL2411P) at a refresh rate of 60 Hz. Audible stimuli were presented diotically through a set of open-back circumnaural headphone (Sennheiser HD650) at a moderate listening level. During the experiment, participants were seated comfortably in a dark, sound-attenuated, and electrically shielded room (Industrial Acoustics Company) approximately 70 cm from the video screen.

### Procedure

The experiment was divided into two parts: rehearsal and testing. In the rehearsal phase, five videos were selected for a participant to learn (randomized across participants). Each of these videos was presented to the participant ten times in a random order with its accompanying audio soundtrack. Participants were instructed to watch and listen attentively and were aware that they would be tested on silent versions of the video after rehearsal. The rehearsal was self-paced with participants advancing to the next trial with a button press. After rehearsal, participants were offered a 10-minute break before testing.

During the testing phase, we collected EEG from participants as they were presented the same five videos encountered during testing plus five new videos (also randomized across participants). The speech audio was stripped from each video (the click was left intact). These silent videos, five “rehearsed” and five “novel,” were repeated four times each and their order was randomized. Before the presentation of each silent rehearsed video, the same video was played with its audio intact to refresh participants’ rehearsal. Prior to each trial, participants were given a target word to detect in the upcoming speech. The target word (see Table 1 for a random selection) was unique across repetitions of a video. Participants were instructed to press the space bar as soon as they perceived the speaker uttering the target. Presses within 2 seconds of target-word utterances were considered a hit (H) and all other presses were recorded as false alarms (FA). We calculated a sensitivity index from hit and false alarm rate using the formula:

**Table 1.** Random selection of target words

|  |  |  |  |
| --- | --- | --- | --- |
| organization | decade | political | healthcare |
| influence | congress | hospital | foreign |
| cost | special | citizen | administration |
| people | plan | funding | issue |
| economy | stability | profit | lower |

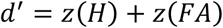

Immediately following each trial, participants were asked to rate the subjective intelligibility of the speech on a ten-point scale, based on the percentage of words they felt they understood.

### EEG Acquisition and Preprocessing

Continuous EEG data were acquired (Figure 1a) using an ActiveTwo system (BioSemi) from 128 scalp electrodes + 2 reference electrodes adhered to the skin over the mastoids. The data were low pass filtered online below 134 Hz and digitized at a rate of 512 Hz. Triggers were sent by an Arduino Uno microcontroller which detected the audio click at the start of each soundtrack to indicate the start of each trial. Subsequent pre-processing was conducted offline in MATLAB: the data were imported and re-referenced to the average reference, bandpass filtered between 0.3 Hz and 15 Hz, and noisy channels were interpolated. To identify channels with excessive noise, the time series were visually inspected in Cartool [75], and the standard deviation of each channel was compared with that of the surrounding channels in MATLAB. Channels contaminated by noise (twice the average standard deviation of the surrounding electrodes) were replaced by spline-interpolating the remaining clean channels with weightings based on their relative scalp location in EEGLAB [76]. Finally, the EEG was epoched and downsampled to 64 Hz.

### Speech Representations

To investigate the links between visual speech and low frequency EEG, we quantified a set of features intended to capture the information in redundant and complementary cues [34] (Figure 1b). As a measure of the redundant visual cue, we extract lip movements (L) using the Face Tracker’s Detailed Features mode in Adobe After Effects, which provided x- and y-coordinates of a series of facial landmarks (namely the center of both lips, corners of the mouth, nose, chin, and both cheeks). A time-series of Euclidean distances between the center of the upper and lower lips defined vertical mouth movements while the distances between the left and right corners of the mouth defined horizontal mouth movements. To account for occasional changes in the camera position we divided vertical lip distance obtained in each frame by the average nose-to-chin distance during the current camera position. Horizontal movements were similarly normalized by the average cheek-to-cheek distance. To capture complementary visual information, we labeled the occurrence of visemes (V) during the speech. Visemes are defined in terms of groupings of phonemes based on perceptual similarity [72,73]. To derive this visemic representation for our videos, we first obtained their phonemic representation [after 43], and mapped each of the 39 phonemes to one of 12 visemes based on the correspondence defined by Auer and Bernstein [37]. There are fewer visemes than phonemes because of the difficulty of distinguishing, for example, a /p/ from a /b/ visually. We also quantified a control measure, motion flow (M), to capture general visual motion in the scene. Briefly [after 38], for each frame, a matrix of motion vectors was calculated using an “Adaptive Rood Pattern Search” block matching algorithm [77]. Motion flow was the sum of all motion vector lengths in each frame [78]. This unweighted sum theoretically could contain both “global motion” (attributable to locomotion or gaze shift) and “local motion” (attributable to external movements), we assume the contribution to this feature from “global motion” was minimal due to the camera remaining stationary relative to the speaker aside from occasional cuts. This signal was then upsampled from 30 Hz to 64 Hz to match the rate of the EEG data. Continuous features motion flow and lip movements were normalized by taking their z-score along the time dimension. Visemes were represented as discrete binary variables indicating the presence (1, blue in Figure 1b) or absence (0) of each viseme at a given time point.

### Temporal Response Function Analysis

In order to relate continuous EEG to the speech representations introduced above, we used a ridge regression analysis that describes the mapping from a stimulus feature of interest to the EEG response in each electrode. This mapping, known as a Temporal Response Function (TRF), was computed and cross-validated using a custom-built toolbox in MATLAB [79]. Because the effect of an event in the environment is not evident in brain activity until after the even and lasts for several hundred milliseconds, we compute the TRF across a series of time lags. To normalize TRFs for comparison across stimulus features, we selected a range of time lags (0-300ms; Figure 1c, shaded regions) that contained the major components of the TRF across features. We constructed a family of TRF models derived from single features or combinations of features and measured their ability to predict new EEG data to estimate the representation of each stimulus feature, or combination of features (Figure 1d). prediction accuracy was measured as the pearson’s correlations between the observed and predicted EEG signals. Where applicable, TRF prediction accuracies were tested against a null distribution generated by iteratively fitting TRFs on shuffled stimulus/response pairings and then evaluating on matched data.

Due to inherent correlations between stimulus features, it was necessary to isolate the effects of single features. To this end, before fitting and evaluating TRFs as described above, we partialed out the contributions of other features from the EEG using the following procedure. First, TRFs were fit to the to-be-partialed feature(s). These TRFs were then used to generate EEG predictions which were subtracted from the original EEG signal. The resultant residual EEG was then used in place of the original EEG in further TRF analyses as described above (Figure 1c, grey arrows). To quantify how well the partialing procedure worked, we measured how well a feature (M, L, or V) can predict residual EEG after partialing (pa_res_; data for V are shown in figure 4b, bottom row). Although the partialing procedure did not reduce prediction accuracy to the level of the noise floor (pa_res,null_; all T(15) > 2.40; all p < 0.03), it does substantially reduce prediction accuracy (Table 2) compared to predicting the original EEG (pa_full_; values from data shown in figure 4a) we calculated this reduction as:

**Table 2.**
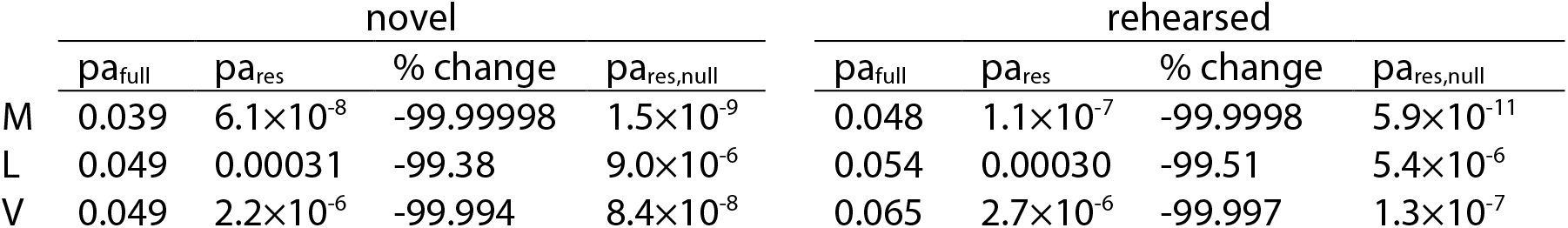
Mean prediction accuracy across occipital channels before and after partialing procedure, the respective percent change, and the noise floor.

|  | novel |  |  |  | rehearsed |  |  |  |
| --- | --- | --- | --- | --- | --- | --- | --- | --- |
| | $pa_{full}$ | $pa_{res}$ | % change | $pa_{res,null}$ | $pa_{full}$ | $pa_{res}$ | % change | $pa_{res,null}$ |
| M | 0.039 | $6.1 \times 10^{-8}$ | -99.99998 | $1.5 \times 10^{-9}$ | 0.048 | $1.1 \times 10^{-7}$ | -99.9998 | $5.9 \times 10^{-11}$ |
| L | 0.049 | 0.00031 | -99.38 | $9.0 \times 10^{-6}$ | 0.054 | 0.00030 | -99.51 | $5.4 \times 10^{-6}$ |
| V | 0.049 | $2.2 \times 10^{-6}$ | -99.994 | $8.4 \times 10^{-8}$ | 0.065 | $2.7 \times 10^{-6}$ | -99.997 | $1.3 \times 10^{-7}$ |

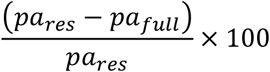

## Acknowledgements

The work and author ARN were supported by NIH grant DC016297 to ECL. Additional support was provided by a Science Foundation Ireland Career Development Award (CDA/15/3316) to ECL. We would like to give a special thanks to Lauren Szymula for help with data collection and to Dr. David Simon for numerous discussions that added substantial value to the manuscript.

## Competing Interests

The authors declare that they have no competing interests, financial or otherwise.

## Notes

### Competing Interest Statement

The authors have declared no competing interest.

## References

1. Sumby WH, Pollack I: Visual Contribution to Speech Intelligibility in Noise. J Acoust Soc Am 1954, 26:212–215.

2. Erber NP: Interaction of audition and vision in the recognition of oral speech stimuli. J Speech Lang Hear Res 1969, 12:423.

3. Zion Golumbic E, Cogan GB, Schroeder CE, Poeppel D: Visual input enhances selective speech envelope tracking in auditory cortex at a “Cocktail Party.– J Neurosci 2013, 33:1417–1426.

4. O’Sullivan AE, Lim CY, Lalor EC: Look at me when I’m talking to you: Selective attention at a multisensory cocktail party can be decoded using stimulus reconstruction and alpha power modulations. Eur J Neurosci 2019, 50.

5. Crosse MJ, Butler JS, Lalor EC: Congruent Visual Speech Enhances Cortical Entrainment to Continuous Auditory Speech in Noise-Free Conditions. J Neurosci 2015, 35:14195–14204.

6. Chandrasekaran C, Trubanova A, Stillittano S, Caplier A, Ghazanfar AA: The natural statistics of audiovisual speech. PLoSComput Biol 2009, 5.

7. Plass J, Brang D, Suzuki S, Grabowecky M: Vision perceptually restores auditory spectral dynamics in speech. Proc Natl Acad Sci US A 2020, 117:16920–16927.

8. Mcgurk H, Macdonald J: Hearing lips and seeing voices. Nature 1976, 264:746–748.

9. Andersson U, Lyxell B, Rönnberg J, Spens K: Cognitive Correlates of Visual Speech Understanding in Hearing-Impaired Individuals. J Deaf Stud Deaf Educ 2001, 6:103–116.

10. Sams M, Aulanko R, Hämäläinen M, Hari R, Lounasmaa O V., Lu ST, Simola J: Seeing speech: visual information from lip movements modifies activity in the human auditory cortex. Neurosci Lett 1991, 127:141–145.

11. Calvert GA, Bullmore ET, Brammer MJ, Campbell R, Williams SCR, McGuire PK, Woodruff PWR, Iversen SD, David AS: Activation of auditory cortex during silent lipreading. Science (80-) 1997, 276:593–596.

12. Ludman CN, Summerfield AQ, Hall D, Elliott M, Foster J, Hykin JL, Bowtell R, Morris PG: Lip-reading ability and patterns of cortical activation studied using fMRI. Br J Audiol 2000, 34:225–230.

13. Pekkola J, Ojanen V, Autti T, Jääskeläinen IP, Möttönen R, Tarkiainen A, Sams M: Primary auditory cortex activation by visual speech: An fMRI study at 3 T. Neuroreport2005, doi:10.1097/00001756-200502080-00010.

14. Besle J, Fischer C, Bidet-Caulet A, Lecaignard F, Bertrand O, Giard MH: Visual activation and audiovisual interactions in the auditory cortex during speech perception: Intracranial recordings in humans. J Neurosci 2008, 28:14301–14310.

15. Bourguignon M, Baart M, Kapnoula EC, Molinaro N: Lip-reading enables the brain to synthesize auditory features of unknown silent speech. J Neurosci 2020, 40:1053–1065.

16. Luo H, Poeppel D: Phase Patterns of Neuronal Responses Reliably Discriminate Speech in Human Auditory Cortex. Neuron 2007, 54:1001–1010.

17. Etard O, Reichenbach T: Neural Speech Tracking in the Theta and in the Delta Frequency Band Differentially Encode Clarity and Comprehension of Speech in Noise. J Neurosci 2019, 39:5750–5759.

18. Peelle JE, Gross J, Davis MH: Phase-Locked Responses to Speech in Human Auditory Cortex are Enhanced During Comprehension. Cereb Cortex 2013, 23:1378–1387.

19. Lalor EC, Power AJ, Reilly RB, Foxe JJ: Resolving Precise Temporal Processing Properties of the Auditory System Using Continuous Stimuli. J Neurophysiol 2009, doi:10.1152/jn.90896.2008.

20. Prinsloo KD, Lalor EC: General auditory and speech-specific contributions to cortical envelope tracking revealed using auditory chimeras. bioRxiv2020, doi:10.1101/2020.10.21.348557.

21. Schroeder CE, Foxe J: Multisensory contributions to low-level, “unisensory” processing. Curr Opin Neurobiol 2005, 15:454–458.

22. Ghazanfar AA, Schroeder CE: Is neocortex essentially multisensory? Trends Cogn Sci 2006, 10:278–285.

23. Kayser C, Petkov CI, Logothetis NK: Visual modulation of neurons in auditory cortex. Cereb Cortex 2008, 18:1560–1574.

24. Lemus L, Hernández A, Luna R, Zainos A, Romo R: Do sensory cortices process more than one sensory modality during perceptual judgments? Neuron 2010, 67:335–348.

25. Cohen L, Rothschild G, Mizrahi A: Multisensory integration of natural odors and sounds in the auditory cortex. Neuron 2011, 72:357–369.

26. Parise C V., Ernst MO: Correlation detection as a general mechanism for multisensory integration. Nat Commun 2016, 7:364.

27. Grant KW, Seitz PFP: The use of visible speech cues for improving auditory detection of spoken sentences. J Acoust Soc Am 2000, 108:1197–1208.

28. Maddox RK, Atilgan H, Bizley JK, Lee AK: Auditory selective attention is enhanced by a task-irrelevant temporally coherent visual stimulus in human listeners. Elife 2015, 2015:1–11.

29. Hershenson M: Reaction time as a measure of intersensory facilitation. J Exp Psychol 1962, 63:289–293.

30. Colonius H, Diederich A: Multisensory Interaction in Saccadic Reaction Time: A Time-Window-of-Integration Model. J Cogn Neurosci 2004, 16:1000–1009.

31. Frassinetti F, Bolognini N, Làdavas E: Enhancement of visual perception by crossmodal visuo-auditory interaction. Exp Brain Res 2002, 147:332–343.

32. Metzger BA, Magnotti JF, Wang Z, Nesbitt E, Karas PJ, Yoshor D, Beauchamp MS: Responses to visual speech in human posterior superior temporal gyrus examined with iEEG deconvolution. J Neurosci 2020, 40:6938–6948.

33. Bernstein LE, Liebenthal E: Neural pathways for visual speech perception. Front Neurosci 2014, 8:386.

34. Campbell R: The processing of audio-visual speech: empirical and neural bases. Philos Trans R Soc B Biol Sci 2008, 363:1001–1010.

35. Peelle JE, Sommers MS: Prediction and constraint in audiovisual speech perception. Cortex 2015, 68:169–181.

36. Dodd B: The role of vision in the perception of speech. Perception 1977, 6:31–40.

37. Auer ET, Bernstein LE: Speechreading and the structure of the lexicon: Computationally modeling the effects of reduced phonetic distinctiveness on lexical uniqueness. J Acoust Soc Am 1997, 102:3704–3710.

38. o GM, Lalor EC: Visual Cortical Entrainment to Motion and Categorical Speech Features during Silent Lipreading. Front Hum Neurosci 2017, 10:679.

39. Files BT, Auer ET, Bernstein LE: The visual mismatch negativity elicited with visual speech stimuli. Front Hum Neurosci 2013, 7:371.

40. Hauswald A, Lithari C, Collignon O, Leonardelli E, Weisz N: A Visual Cortical Network for Deriving Phonological Information from Intelligible Lip Movements. Curr Biol 2018, 28:1453-1459.e3.

41. Ozker M, Yoshor D, Beauchamp MS: Frontal cortex selects representation of the talker’s mouth to aid in speech perception. Elife 2018, 7.

42. Mégevand P, Mercier MR, Groppe DM, Zion Golumbic E, Mesgarani N, Beauchamp MS, Schroeder CE, Mehta AD: Crossmodal phase reset and evoked responses provide complementary mechanisms for the influence of visual speech in auditory cortex. J Neurosci 2020, doi:10.1523/JNEUROSCI.0555-20.2020.

43. Low-frequency cortical entrainment to speech reflects phoneme-level processing. Curr Biol 2015, 25:2457–2465.

44. Bednar A, Lalor EC: Where is the cocktail party? Decoding locations of attended and unattended moving sound sources using EEG. Neuroimage 2020, 205:116283.

45. Nunez-Elizalde AO, Huth AG, Gallant JL: Voxelwise encoding models with non-spherical multivariate normal priors. Neuroimage 2019, 197:482–492.

46. Howard MF, Poeppel D: Discrimination of Speech Stimuli Based on Neuronal Response Phase Patterns Depends on Acoustics But Not Comprehension. J Neurophysiol 2010, 104:2500–2511.

47. Smith ZM, Delgutte B, Oxenham AJ: Chimaeric sounds reveal dichotomies in auditory perception. Nature 2002, 416:87–90.

48. Ross LA, Saint-Amour D, Leavitt VM, Javitt DC, Foxe JJ: Do you see what I am saying? Exploring visual enhancement of speech comprehension in noisy environments. Cereb Cortex 2007, 17:1147–1153.

49. Bernstein LE, Demorest ME, Tucker PE: Speech perception without hearing. Percept Psychophys 2000, 62:233–252.

50. Bernstein LE, Jiang J, Pantazis D, Lu Z-L, Joshi A: Visual phonetic processing localized using speech and nonspeech face gestures in video and point-light displays. Hum Brain Mapp 2011, 32:1660–1676.

51. Calvert GA, Campbell R: Reading speech from still and moving faces: The neural substrates of visible speech. J Cogn Neurosci 2003, 15:57–70.

52. Nidiffer AR, Diederich A, Ramachandran R, Wallace MT: Multisensory perception reflects individual differences in processing temporal correlations. Sci Rep 2018, 8.

53. Parise C V., Spence C, Ernst MO: When correlation implies causation in multisensory integration. Curr Biol 2012, 22:46–49.

54. Nozaradan S, Peretz I, Mouraux A: Steady-state evoked potentials as an index of multisensory temporal binding. Neuroimage 2012, 60:21–28.

55. Parise C V, Harrar V, Ernst MO, Spence C: Cross-correlation between Auditory and Visual Signals Promotes Multisensory Integration. Multisens Res 2013, 26:1–10.

56. Bizley JK, Maddox RK, Lee AK: Defining Auditory-Visual Objects: Behavioral Tests and Physiological Mechanisms. Trends Neurosci 2016, 39:74–85.

57. Nidiffer AR, Ramachandran R, Wallace MT: Multisensory binding is driven by the strength of stimulus correlation. PsyArXiv 2020, doi:10.31234/osf.io/vzbmw.

58. Park H, Kayser C, Thut G, Gross J: -frequency brain oscillations to facilitate speech intelligibility. Elife 2016, 5:e14521.

59. Atilgan H, Town SM, Wood KC, Jones GP, Maddox RK, Lee AKC, Bizley JK: Integration of Visual Information in Auditory Cortex Promotes Auditory Scene Analysis through Multisensory Binding. Neuron 2018, 97:640-655.e4.

60. O’Sullivan A Crosse M, Di Liberto G, de Cheveigné A, Lalor E: Neurophysiological indices of audiovisual speech integration are enhanced at the phonetic level for speech in noise. bioRxiv 2020, doi:10.1101/2020.04.18.048124.

61. Blank I, Balewski Z, Mahowald K, Fedorenko E: Syntactic processing is distributed across the language system. Neuroimage 2016, 127:307–323.

62. Broderick MP, Anderson AJ, Lalor EC: Semantic Context Enhances the Early Auditory Encoding of Natural Speech. J Neurosci 2019, doi:10.1523/JNEUROSCI.0584-19.2019.

63. Anderson AJ, Binder JR, Fernandino L, Humphries CJ, Conant LL, Raizada RDS, Lin F, Lalor EC: An Integrated Neural Decoder of Linguistic and Experiential Meaning. J Neurosci 2019, 39:8969–8987.

64. Cappe C, Barone P: Heteromodal connections supporting multisensory integration at low levels of cortical processing in the monkey. Eur J Neurosci 2005, 22:2886–2902.

65. Bizley JK, Nodal FR, Bajo VM, Nelken I, King AJ: Physiological and anatomical evidence for multisensory interactions in auditory cortex. Cereb Cortex 2007, 17:2172–2189.

66. Falchier A, Schroeder CE, Hackett TA, Lakatos P, Nascimento-Silva S, Ulbert I, Karmos G, Smiley JF: Projection from Visual Areas V2 and Prostriata to Caudal Auditory Cortex in the Monkey. Cereb Cortex 2010, 20:1529–1538.

67. Nath AR, Beauchamp MS: Dynamic changes in superior temporal sulcus connectivity during perception of noisy audiovisual speech. J Neurosci 2011, 31:1704–1714.

68. Bisley JW: The neural basis of visual attention. J Physiol 2011, 589:49–57.

69. Shinn-Cunningham BG: Object-based auditory and visual attention. Trends Cogn Sci 2008, 12:182–186.

70. Desimone R, Duncan J: Neural Mechanisms of Selective Visual Attention. Annu Rev Neurosci 1995, 18:193–222.

71. Daube C, Ince RAA, Gross J: Simple Acoustic Features Can Explain Phoneme-Based Predictions of Cortical Responses to Speech Article Simple Acoustic Features Can Explain Phoneme-Based Predictions of Cortical Responses to Speech. Curr Biol 2019, 29:1924-1937.e9.

72. Woodward MF, Barber CG: Phoneme Perception in Lipreading. J Speech Hear Res 1960, 3:212–222.

73. Fisher CG: Confusions among visually perceived consonants. J Speech Hear Res 1968, 11:796–804.

74. en Oever S, Schroeder CE, Poeppel D, van Atteveldt N, Zion-Golumbic E: Rhythmicity and cross-modal temporal cues facilitate detection. Neuropsychologia 2014, 63:43–50.

75. Brunet D, Murray MM, Michel CM: Spatiotemporal analysis of multichannel EEG: CARTOOL. Comput Intell Neurosci 2011, 2011.

76. Delorme A, Makeig S: EEGLAB: An open source toolbox for analysis of single-trial EEG dynamics including independent component analysis. J Neurosci Methods 2004, 134:9–21.

77. Barjatya A: Block matching algorithms for motion estimation. IEEE Trans Evol Comput 2004, 8:225–239.

78. Bartels A, Zeki S, Logothetis NK: Natural vision reveals regional specialization to local motion and to contrast-invariant, global flow in the human brain. Cereb Cortex 2008, 18:705–717.

79. Crosse MJ, Di Liberto GM, Bednar A, Lalor EC: The Multivariate Temporal Response Function (mTRF) Toolbox: A MATLAB Toolbox for Relating Neural Signals to Continuous Stimuli. Front Hum Neurosci 2016, 10:604.

